# Chemical inhibition of prolyl hydroxylases impairs angiogenic competence of human vascular endothelium through metabolic reprogramming

**DOI:** 10.1101/2022.03.25.485858

**Authors:** Ratnakar Tiwari, Prashant V. Bommi, Peng Gao, Matthew J. Schipma, Yalu Zhou, Susan E. Quaggin, Navdeep S. Chandel, Pinelopi P. Kapitsinou

**Affiliations:** Feinberg Cardiovascular Research Institute, Northwestern University Feinberg School of Medicine, Chicago, IL; Division of Nephrology & Hypertension, Northwestern University Feinberg School of Medicine, Chicago, IL, USA; Department of Medicine and Robert H. Lurie Cancer Center, Northwestern University Feinberg School of Medicine, Chicago, IL; Department of Biochemistry and Molecular Genetics, Feinberg School of Medicine, Northwestern University, Chicago, IL

**Keywords:** PHDs, HIFs, endothelium, metabolism, angiogenesis

## Abstract

Endothelial cell (EC) metabolism has emerged as a driver of angiogenesis. While hypoxia inactivates prolyl-4 hydroxylase domain containing proteins 1-3 (PHD1-3) and stabilizes hypoxia inducible factors (HIFs) stimulating angiogenesis, the effects of PHDs on EC functions remain unclear. Here, we investigated the impact of PHD inhibition by dimethyloxalylglycine (DMOG) on angiogenic competence and metabolism of human vascular ECs. PHD inhibition reduced EC proliferation, migration, and tube formation capacities. Furthermore, transcriptomic and metabolomic analyses revealed an unfavorable metabolic reprogramming for angiogenesis following treatment with DMOG. Despite the induction of glycolytic genes and high levels of lactate, multiple genes encoding sub-units of mitochondrial complex I were suppressed with concurrent decline in nicotinamide adenine dinucleotide (NAD^+^) levels. Importantly, defective EC migration due to DMOG could be partially restored by augmenting NAD^+^ levels. Combined, our data provide metabolic insights into the mechanism by which chemical PHD inhibition impairs angiogenic competence of human vascular ECs.

## INTRODUCTION

Endothelial cells (ECs) line the blood vessels and play a critical role in providing oxygen and nutrients to all tissues in multicellular organisms. In adult tissues, normal ECs remain in quiescence, but injury can induce endothelial activation and angiogenesis, allowing delivery of oxygen and nutrients to hypoxic tissues. Recently, metabolism has emerged as a key regulator of angiogenesis in parallel to well-established angiogenic growth factors (1). ECs are highly glycolytic and 6-phosphofructo-2-kinase/fructose-2,6-bisphosphatase 3 (PFKFB3)-driven glycolysis controls vessel sprouting (2). Furthermore, fatty acid oxidation promotes EC proliferation through DNA synthesis (3), while glutamine metabolism replenishes the mitochondrial tricarboxylic acid (TCA) cycle and maintains EC sprouting (4, 5). Finally, mitochondrial respiratory chain complex III was recently shown to be required for EC proliferation by sustaining amino acid availability (6).

Since ECs reside in direct contact with oxygen, it is not surprising that they are equipped with various O_2_ sensing mechanisms able to regulate EC function with critical impact in surrounding tissues. A key pathway mediating hypoxia signaling is oxygen-dependent degradation of hypoxia-inducible factors (HIFs), heterodimeric transcription factors made up of an oxygen-labile α-subunit (HIF-α) and a stable β-subunit (HIF-β). Under normoxia, HIF-α undergoes proline hydroxylation, a post-translational modification mediated by 2-oxoglutarate, iron dioxygenases called prolyl-4-hydroxylase domain (PHD) proteins 1-3 (7, 8). Proline hydroxylation allows recognition and ubiquitination of HIF-α by the tumor suppressor protein von Hippel-Lindau (pVHL) followed by proteasomal degradation (9). Conversely, in hypoxia, PHDs are inactivated and HIF-α subunits are stabilized and induce the expression of genes promoting adaptation to low O_2_ such as metabolic and angiogenic genes (10, 11). It is now well established that HIFs control angiogenesis in embryonic vascular development and disease settings such as cancer and vascular diseases (12).

Besides sensing oxygen and metabolic stimuli such as amino acid, α-ketoglutarate, succinate, or fumarate (13-15), PHDs have emerged as regulators of metabolism. For instance, loss of PHD1 reduces mitochondrial respiration in muscle (16) and enhances glucose flux through the oxidative pentose phosphate pathway in neurons (17). PHD2 deficiency induces anaerobic glycolysis in macrophages (18), while PHD3 loss enhances mitochondrial fat metabolism in muscle (19). Nevertheless, the impact of PHD inhibition in EC metabolism remains poorly defined. Given the importance of endothelial oxygen sensing and metabolism for vascular growth and function, we investigated the angiogenic competence and metabolic profile of EC subjected to chemical PHD inhibition.

## RESULTS

### Chemical inhibition of PHDs impairs EC proliferation, migration, and tube formation

In response to angiogenic stimuli, EC proliferation and migration are prerequisite processes for the formation of blood vessels (20-22). To define the impact of PHD inhibition on these essential angiogenic processes, we exposed human primary pulmonary artery endothelial cells (HPAEC) to dimethyloxalylglycine (DMOG), a cell permeable competitive PHD inhibitor. As expected, treatment with DMOG (1 mM) for 24 h led to nuclear stabilization of HIF-1α and HIF-2α as shown by immunofluorescence staining and immunoblot analysis (Supplemental Fig. 1A and B) without affecting cell viability based on Annexin V/ Propidium Iodide (PI) staining (Supplemental Fig. 1C). To assess the effect of chemical PHD inhibition on HPAEC proliferation, we first used the MTT colorimetric assay, which is based on cell metabolic activity. We observed a ∼40% reduction in proliferation of EC treated with DMOG compared to controls (Fig. 1A). This effect was validated by BrdU immunofluorescence labeling, which showed reduction in number of BrdU positive nuclei per high-power field (hpf) by 84% in DMOG-exposed cells compared to vehicle-treated cells (P < 0.001) (Fig. 1B). Furthermore, flow cytometric cell cycle analysis by DNA staining with propidium iodide indicated suppression of cell cycle progression in DMOG-treated cells (Fig. 1C). Specifically, treatment with DMOG increased by ∼6% the proportion of cells in G0/G1 phase (P <0.01) and reduced by ∼48% the proportion of cells in S phase (P <0.0001). We also observed a reduction of cells in G2/M phase, which though did not reach statistical significance (P=0.057).

**Figure 1.**
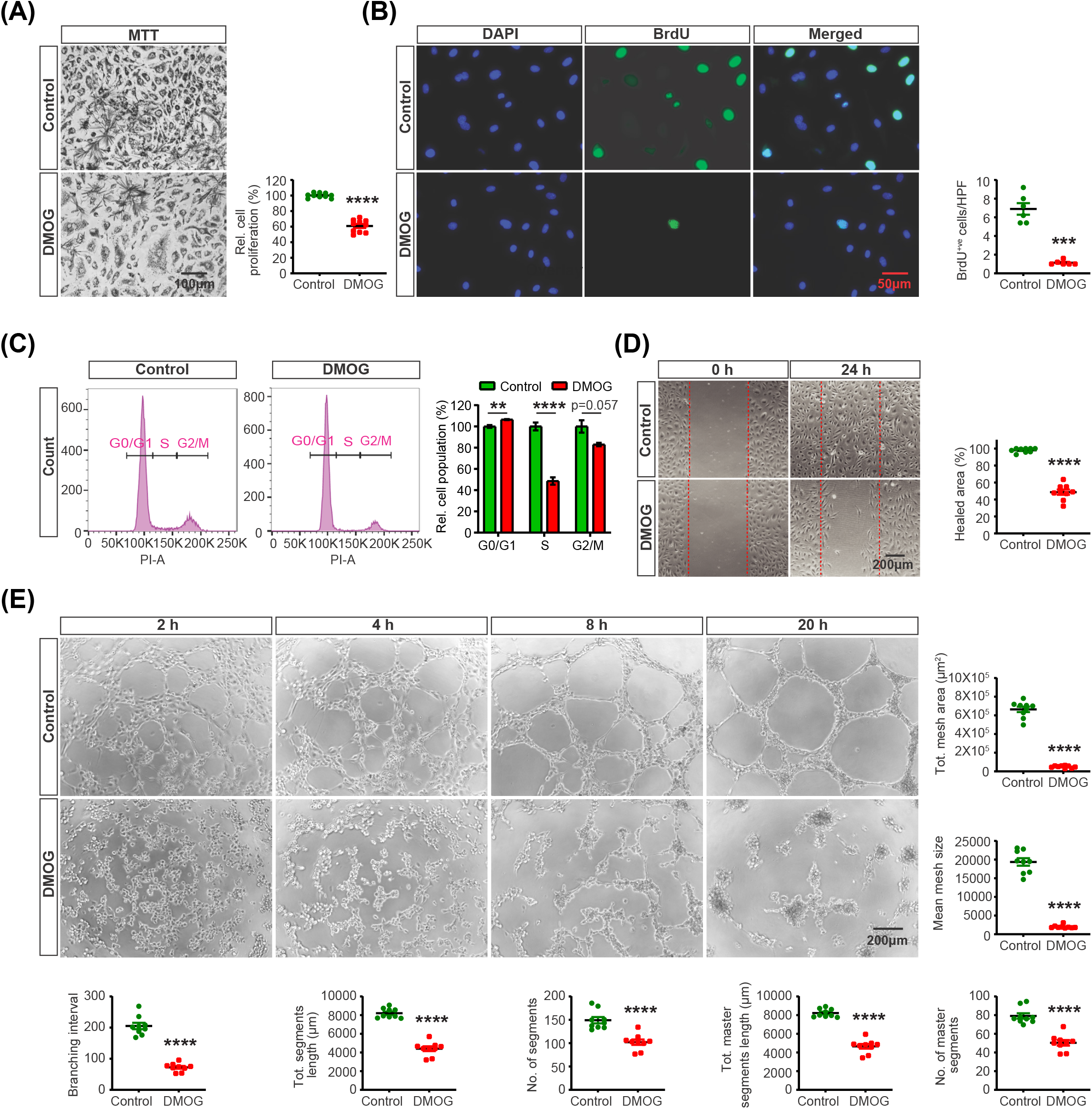
Prolyl hydroxylase inhibition by DMOG suppresses endothelial cell proliferation, migration, and tube formation. (**A**) Representative bright field images of formazan crystal formed after 3 h incubation of MTT with vehicle or DMOG-treated (1 mM) HPAEC. Right graph shows relative HPAEC proliferation assessed by MTT assay. (**B**) Representative images of BrdU immunostaining. Right graph shows semi-quantitative analysis of BrdU positive cells/hpf. (**C**) Representative histograms of cell cycle analysis for control and DMOG-treated cells. Right side graph demonstrates relative percentages of cell populations in G0/G1, S and G2/M cell cycle phases. (**D**) Representative images of 2D scratch wound assay of control and DMOG-treated cells and semi-quantitative analysis of healed area after 24 h. (**E**) Representative images of tubes formed at indicated time points in control and DMOG-treated cells and semi-quantitative analysis of different parameters at 20 h time point. Data are pooled from 3 independent experiments and represented as mean ± SEM. Statistics were determined by two-tailed t-test. **, P<0.01; ****, P< 0.0001; ns, not statistically significant.

Next, we conducted a 2D scratch-wound assay to determine the effect of DMOG on migratory capacity of HPAEC. As shown in Figure 1D, we found a ∼49% reduction in closure of the wounded gap in DMOG-treated EC compared to vehicle (P <0.0001). In addition, we assessed the effect of DMOG on EC tube formation. Quantitative analysis of several angiogenic parameters including total meshes area and segments length revealed that DMOG significantly disrupted EC tube formation (Fig. 1E). Taken together, our findings show that chemical inhibition of PHDs by DMOG impairs angiogenic competence of HPAEC.

### Chemical inhibition of PHDs induces a broad transcriptional response in HPAEC

We next performed RNA-seq analysis to define the transcriptional response of HPAEC to DMOG-induced PHD inhibition. HPAEC were exposed to 1 mM DMOG or vehicle (0.5% DMSO) for 24 h prior to analysis (4 DMOG and 4 vehicle-treated samples). RNA-seq identified 4,092 differentially expressed protein coding genes (DEGs), of which 1,000 were up-regulated and 3,092 were down-regulated (Supplemental Fig. 2A). Heatmap of the top 50 among the significantly up-regulated and down-regulated genes showed a distinct transcriptional profile following treatment with DMOG (Supplemental Fig. 2B). Hallmark analysis of DEGs showed that the transcriptional response to DMOG treatment mimicked hypoxia, validating activation of the HIF pathway (Fig. 2A). Similarly, HIF-1 signaling pathway was among the top enriched pathways detected by KEGG pathway analysis of DEGs (Fig. 2B). Importantly, Hallmark and KEGG pathways showed significant alteration of genes involved in cellular metabolism (Fig. 2A and B). Specifically, exposure to DMOG significantly induced genes involved in glucose metabolism (Fig. 2C and Supplemental Fig. 2C), as indicated by the significant up-regulation of the glucose transporters *solute carrier family 2 member 1* (*SLC2A1*) and *solute carrier family 2 member 3* (*SLC2A3*) and glycolytic genes *hexokinase 1* (*HK1*), *hexokinase 2* (*HK2*), *6-phosphofructo-2-kinase/fructose-2,6-biphosphatase 3* (*PFKFB3*), *and lactate dehydrogenase A* (*LDHA*). In contrast, DMOG suppressed several genes involved in mitochondrial metabolism (Fig. 2C, D and Supplemental Fig. 2C). Among TCA cycle genes, we observed down-regulation of *isocitrate dehydrogenase (IDH3A), succinyl-CoA ligase* (*SUCLA2)* and *succinate dehydrogenase (SDHC)* by DMOG. Furthermore, genes encoding sub-units of mitochondrial complex I (MC1), which regenerates NAD^+^, were significantly suppressed by DMOG (Fig. 2D and Supplemental Fig. 2C). Similarly, *succinate dehydrogenase complex assembly factor 3 and 4* (*SDHAF3 and SDHAF4*) of mitochondrial complex II were significantly downregulated, whereas no significant alterations were noted in genes encoding sub-units of mitochondrial complexes III, IV and V (Fig. 2D and Supplemental Fig. 2C).

**Figure 2.**
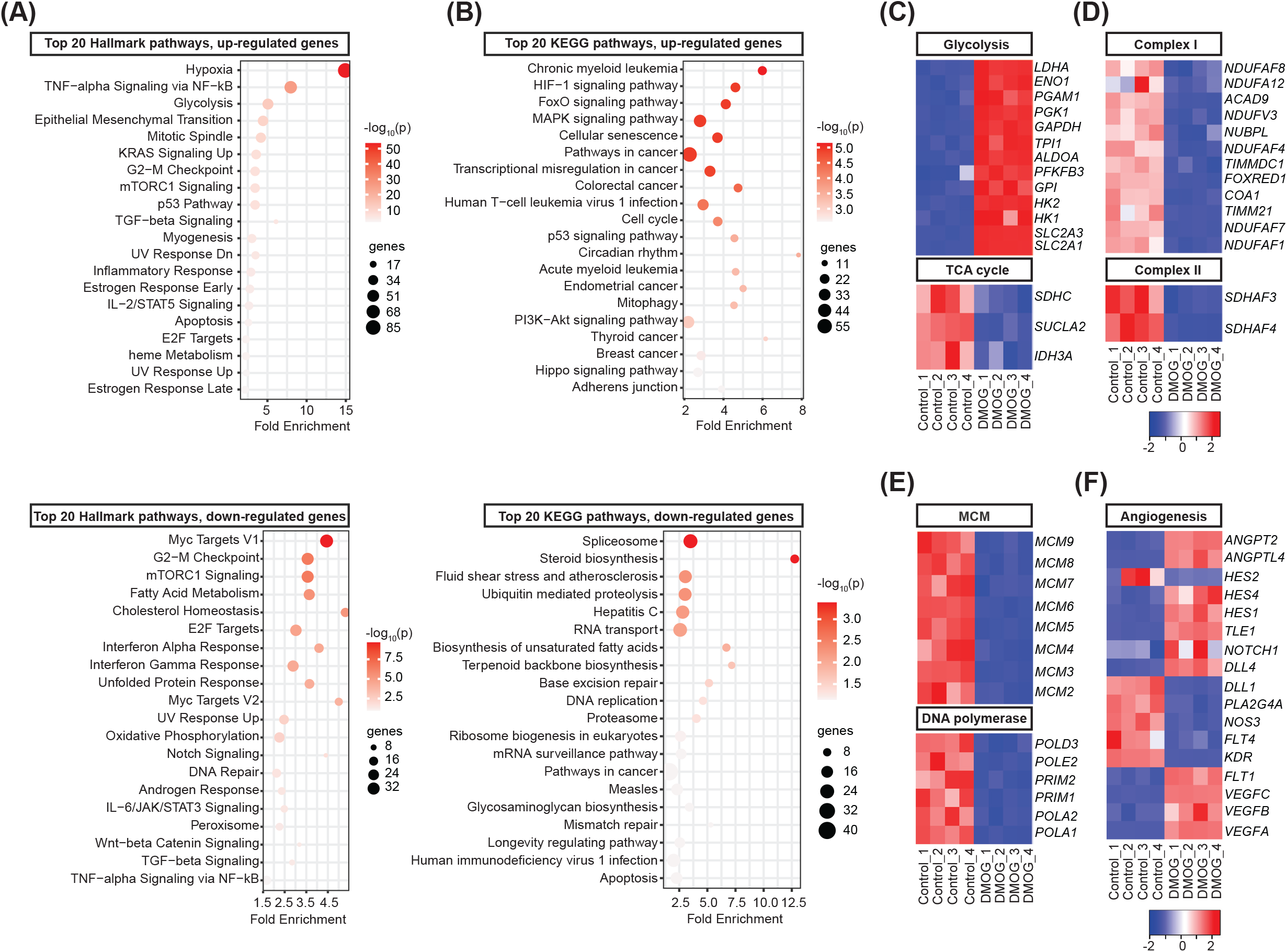
Prolyl hydroxylase inhibition by DMOG mimics transcriptional response to hypoxia, altering the expression of genes involved in metabolism, cell cycle and angiogenesis. Bubble charts for top 20 enriched Hallmark (**A**) and KEGG (**B**) pathways of up-regulated (top) or down-regulated DEGs (bottom) by DMOG. Heat maps of RNA-seq data showing significantly altered genes involved in glycolysis and TCA cycle (**C**), mitochondrial electron transport complex I and II (**D**), cell cycle (**E**) and angiogenesis (**F**). Each column corresponds to a sample and each row corresponds to a specific gene. KEGG, Kyoto Encyclopedia of Genes and Genomes.

Consistent with the reduced proliferation capacity of HPAEC treated with DMOG, cell cycle was identified among the enriched pathways in the transcriptome of DMOG-treated cells. Indeed, hypoxia suppresses genes involved in DNA replication (23) and DMOG reduced the expression of genes encoding DNA-polymerases, primases and microsomal maintenance (MCM) proteins 2-7, components of a DNA helicase required for DNA replication and cell proliferation (24) (Fig. 2E). Furthermore, mRNA levels of classic angiogenic genes were significantly altered by treatment with DMOG (Fig. 2F). Among genes involved in VEGF signaling, *vascular endothelial growth factor A* (*VEGFA*), *vascular endothelial growth factor B* (*VEGFB*), *vascular endothelial growth factor C* (*VEGFC*) and *fms related receptor tyrosine kinase 1* (*FLT1*) were significantly up-regulated, while *kinase insert domain receptor* (*KDR*) was down-regulated in DMOG-treated cells compared to control. Significant alterations were also observed in expression of genes in Notch signaling pathway as indicated by down-regulation of *delta-like protein 1* (*DLL1*) and *hes family bHLH transcription factor 2* (*HES2*) and up-regulation of *delta like canonical Notch ligand 4* (*DLL4*), *notch receptor 1* (*NOTCH1*), *transducin-like enhancer of split 1* (*TLE1*), *hes family bHLH transcription factor 1* (*HES1*) and *hes family bHLH transcription factor 4* (*HES4*) in the setting of DMOG exposure. Finally, among angiopoietins and angiopoietin-like proteins, DMOG induced the mRNAs of *angiopoietin 2 (ANGPT2*) and *angiopoietin-like 4* (*ANGPTL4*) (Fig. 2F). Taken together, PHD inhibition by DMOG induces a broad transcriptional response in HPAEC with significant alterations in the expression of genes involved in metabolism, cell cycle, and angiogenesis.

### Chemical inhibition of PHDs induces metabolic reprogramming in HPAEC

To get insights into how DMOG exposure affects EC metabolism, we assessed the metabolome of cell lysates and media of HPAEC treated with DMOG compared to vehicle (5 DMOG-treated and 5 vehicle-treated samples). Unbiased metabolomic profiling by liquid chromatography/mass spectrometry (LC/MS) and gas chromatography/mass spectrometry (GC/MS) platforms showed significant alterations in 126 metabolites for cell lysates (70 up-regulated and 56 down-regulated) and 45 metabolites for cell media (19 up-regulated and 26 down-regulated) in the setting of DMOG treatment (Supplemental Fig. 3A, Supplemental Table sheet S1 and 2). Biochemical importance plot generated by random forest classification showed key differences in amino acid, nucleotide, lipid and glucose metabolism (Supplemental Fig. 3B and C). Accordingly, metabolites set enrichment analysis using Small Molecule Pathway Database (SMPDB) detected Warburg effect, citric acid, lipid, amino acid, and nucleotide metabolism among the most frequently modified pathways by DMOG (Fig. 3A).

**Figure 3.**
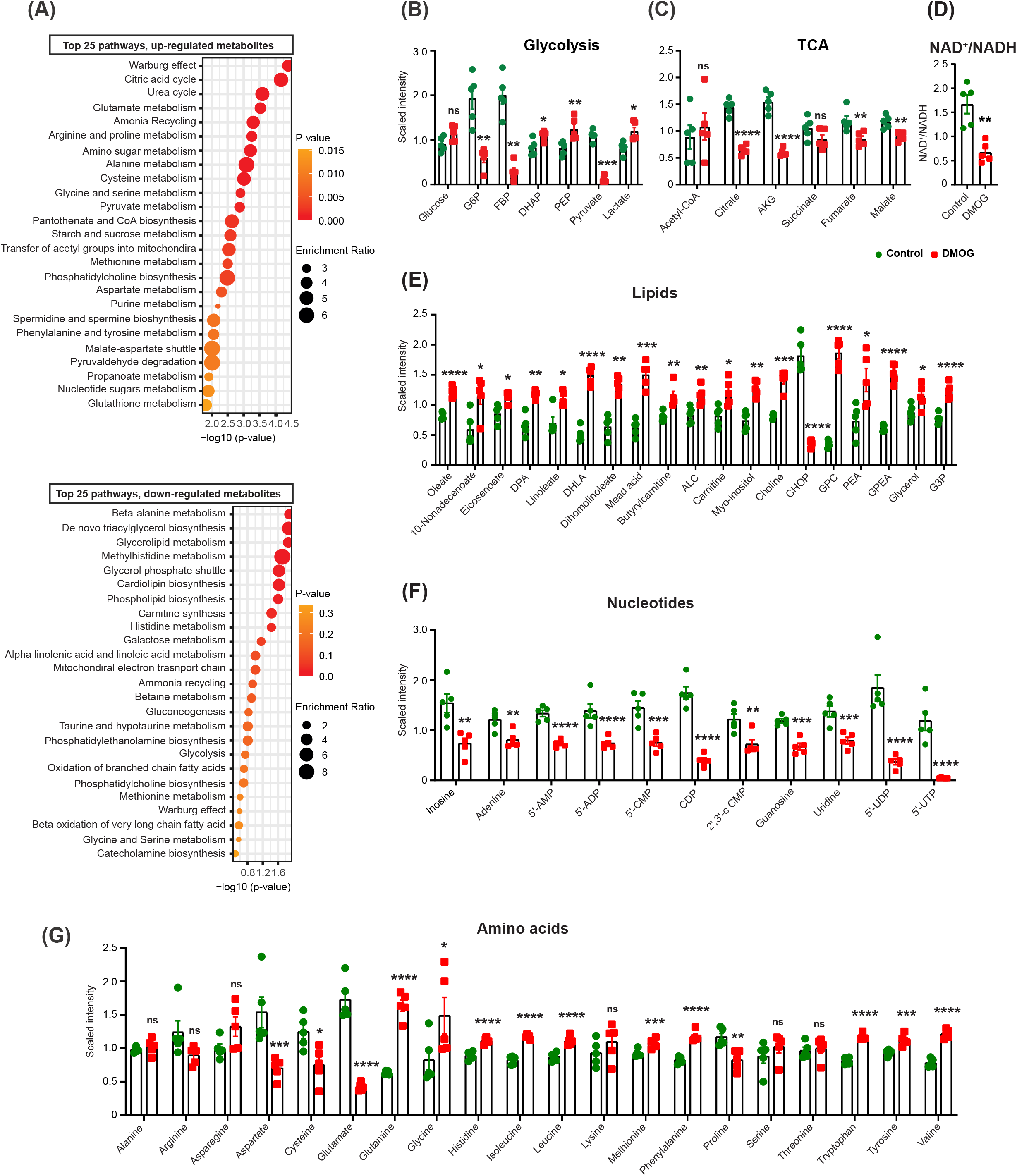
Prolyl hydroxylase inhibition by DMOG alters the endothelial cell metabolome. (**A**) Shown are the top 25 down-regulated (upper graph) and up-regulated (lower graph) metabolic pathways detected by metabolites set enrichment analysis in DMOG-treated cells compared to control. Scaled intensity values indicating relative levels of metabolites related to glycolysis (**B**) and TCA cycle (**C**). (**D**) NAD^+^/NADH ratio in cells treated with vehicle or DMOG. Scaled intensity values indicating relative levels of lipid metabolites (**E**), nucleotides (**F**) and amino acids (**G**). n=5 independent samples per condition. All statistical data are represented as mean ± SEM and statistics were determined by a Welch’s two sample t-test. *, P<0.05; **, P<0.01; ***, P<0.001; ****, P<0.0001; ns, not significant. G6P, glucose-6-phosphate; FBP, fructose 1,6 bisphosphate; DHAP, dihydroxyacetone phosphate; PEP, phosphoenolpyruvate; AKG, alpha-ketoglutarate; DPA, docosapentaenoate; DHLA, dihomolinolenate; ALC, acetylcarnitine; CHOP, choline phosphate; GPC, glycerophosphorylcholine; PEA, phosphoethanolamine; GPEA, glycerylphosphorylethanolamine; G3P, glycerol 3-phosphate; 5’-AMP, adenosine-5’-monophosphate; 5’-ADP, adenosine-5’-diphopshate; 5’-CMP, cytidine 5’-monophosphate; CDP, cytidine diphosphate; 2’,3’-cCMP, cytidine 2’,3’-cyclic monophosphate; 5’-UDP, uridine-5-diphosphate; UTP, uridine 5’-triphosphate.

#### DMOG exposure altered glucose metabolism in HPAEC

DMOG-treated ECs showed significant alterations in several glucose-derived metabolites (Fig. 3B). Compared to vehicle, treatment with DMOG induced significant reduction in levels of 6-carbon glycolytic intermediates, glucose-6-phosphate (G6P) (3.2-fold, P<0.01) and fructose 1,6 bisphosphate (FBP) (7-fold, P<0.01), while significant increase was noted for 3-carbon glycolytic intermediates, dihydroxyacetone phosphate (DHAP) (1.3-fold, P<0.05) and phosphoenolpyruvate (PEP) (1.5-fold, P<0.01). Finally, pyruvate level was reduced by10-fold (P<0.001), while lactate level was increased by 1.46-fold (P<0.05) in DMOG-treated cells.

#### DMOG exposure impairs mitochondrial metabolism

DMOG-treated cells displayed decreased levels for several TCA cycle intermediates compared to the control (Fig. 3C). Specifically, citrate, alpha-ketoglutarate, fumarate, and malate levels were significantly reduced by DMOG. Furthermore, DMOG-treated cells showed significantly reduced NAD^+^/NADH ratio, known to limit carbon entry and flow through the TCA cycle (25) (Fig. 3D). Interestingly, acetyl-CoA levels remained unchanged despite the significant reduction in pyruvate levels, suggesting either reduced carbon-flux of TCA cycle and/or contribution of other pathways in the replenishment of acetyl-CoA pool.

#### DMOG exposure impairs lipid metabolism

Lipids function as structural components of membrane, signaling molecules for proliferation and apoptosis and support TCA cycle (1). After DMOG treatment, significant increase was observed in several lipids including, oleate (1.5-fold, P<0.001) and linoleate (1.5-fold, P<0.05), choline (1.7-old, P<0.001), glycerophosphorylcholine (GPC) (5.3-fold, P<0.001), glycerophosphoethanolamine (GPEA) (2.4 -fold, P<0.001), and glycerol 3-phosphate (G3P) (1.6-fold, P<0.001) (Fig. 3E).

#### DMOG treatment alters nucleotide metabolism

Metabolomic profiling revealed significant alterations in the level of inosine, and several purine and pyrimidine-metabolites (Fig. 3F). Specifically, treatment with DMOG caused significant reduction in inosine, adenine, adenosine-5’-diphopshate (ADP), adenosine-5’-monophosphate (AMP), and guanosine levels. Among pyrimidine metabolites, cytidine 5’-monophosphate (CMP), cytidine diphosphate (CDP), cytidine 2’,3’-cyclic monophosphate (2’,3’-cCMP), uridine, uridine-5-diphosphate (UDP) and uridine 5’-triphosphate (UTP) were significantly reduced by DMOG. The resulting diminished production in nucleotide pool is likely to suppress EC vessel sprouting as reported by Schoors et al. (3).

#### DMOG alters amino acid levels in HPAEC

DMOG exposed HPAEC showed significant changes in the level of amino acids (Fig 3G). Out of 20 amino acids; 13 were significantly altered post DMOG treatment. Specifically, the levels of 4 amino acids (aspartate, glutamate cysteine, and proline) were significantly reduced, while 9 amino acids (glutamine, histidine, isoleucine, leucine, methionine, phenylalanine, tryptophan, tyrosine, and valine) showed significant increase following treatment with DMOG. Importantly, increased glutamine levels in conjunction with the diminished glutamate indicated diminished glutamine utilization, a response that can impair angiogenesis (4, 5). Furthermore, significant reduction was noted in levels of aspartate, known to regulate cell proliferation by supporting the biosynthesis of purines and pyrimidines (26, 27) In summary, these findings suggest that treatment of HPAEC with DMOG induces a switch to anaerobic glycolysis, impairs TCA cycle, and alters broadly amino acid levels with concurrent reduction in purine and pyrimidine levels, metabolic changes, which could lead to reduced HPAEC proliferation and impaired angiogenic competence in the setting of DMOG treatment.

### Supplementation with pyruvate or aspartate fails to rescue DMOG-induced defects in angiogenic competence of HPAEC

Among the metabolic alterations induced by DMOG, we wished to define the role of the observed reductions in pyruvate and aspartate levels. Pyruvate is essential in maintaining carbon flux in TCA cycle and reduced pyruvate levels have been reported to impair mitochondrial metabolism (28-30). Aspartate functions as a precursor for nucleotide synthesis and inhibition of aspartate production from mitochondria or its transport to cytoplasm reduces cell proliferation (27, 31). We therefore examined whether supplementation of these metabolites was sufficient to rescue the angiogenic defects induced by DMOG. Nevertheless, the effects of DMOG on proliferation, migration, and tube formation abilities of HPAEC (Supplemental Fig. 4A, B, C, D and E) remained unchanged by supplementation with methyl pyruvate (2 mM, 1 mM and 0.5 mM) or methyl aspartate (5 mM, 2 mM, 0.5 mM). In the absence of DMOG, methyl pyruvate did not influence endothelial angiogenic competence (Supplemental Fig. 4A,B, and E), while methyl aspartate at high concentrations (5 mM, 2 mM) reduced cell proliferation.

### Supplementation with citrate or nicotinamide riboside partially rescue DMOG-induced defects in endothelial cell migration and tube formation ability

Because the collapse of TCA cycle intermediates in the setting of blocked glutamine metabolism may contribute in DMOG-induced angiogenic defects, we next focused on citrate, an important substrate in TCA cycle, which has been shown to promote cell proliferation (32). Therefore, we supplemented cells with membrane permeable form of citrate (trimethyl citrate) (33). Citrate supplementation (0.5 mM) improved the metabolic activity based MTT assay (Fig. 4A) but had no effect on DMOG induced alterations in cell proliferation and cell cycle progression as indicated by BrdU incorporation assay and flow cytometric analysis, respectively (Fig. 4B and C and Supplemental Fig. 5A). Nevertheless, citrate supplementation partially rescued the DMOG-induced defects on migration and tube formation abilities (Fig. 4D and E) as indicated by a 22% increase in healing area of 2D scratch wound assay (P< 0.0001) (Fig. 4D) and improvement of multiple tube formation-related parameters (Fig. 4E). To investigate the molecular mechanism by which citrate improved the DMOG induced angiogenic defects, we examined the HIF pathway. Notably, treatment with citrate did not alter HIF-1 and HIF-2 protein levels in the setting of DMOG exposure (Supplemental Fig. 5B and C). Because citrate is a metabolic intermediate of reductive carboxylation of glutamine, a process known to recycle NADH to NAD^+^ under impaired mitochondrial metabolism (34), we next examined the impact of citrate supplementation on DMOG-induced reduction in NAD^+^ levels. By performing LC/MS, we found that citrate supplementation in DMOG-treated cells raised NAD^+^ levels by 2.1-fold (Supplemental Fig. 5D). In the absence of DMOG, citrate supplementation did not affect NAD^+^ levels in HPAEC. Furthermore, direct NAD^+^ repletion by adding the NAD^+^ precursor nicotinamide riboside (NR) (200 μM) mirrored the effects of citrate and improved the DMOG-induced defects in migration and tube formation but without affecting EC proliferation (Fig 5A-D). In summary, these findings identify citrate and NAD^+^ depletion as significant metabolic alterations contributing to the impaired EC migration and tube formation capacity in the context of PHD inhibition.

**Figure 4.**
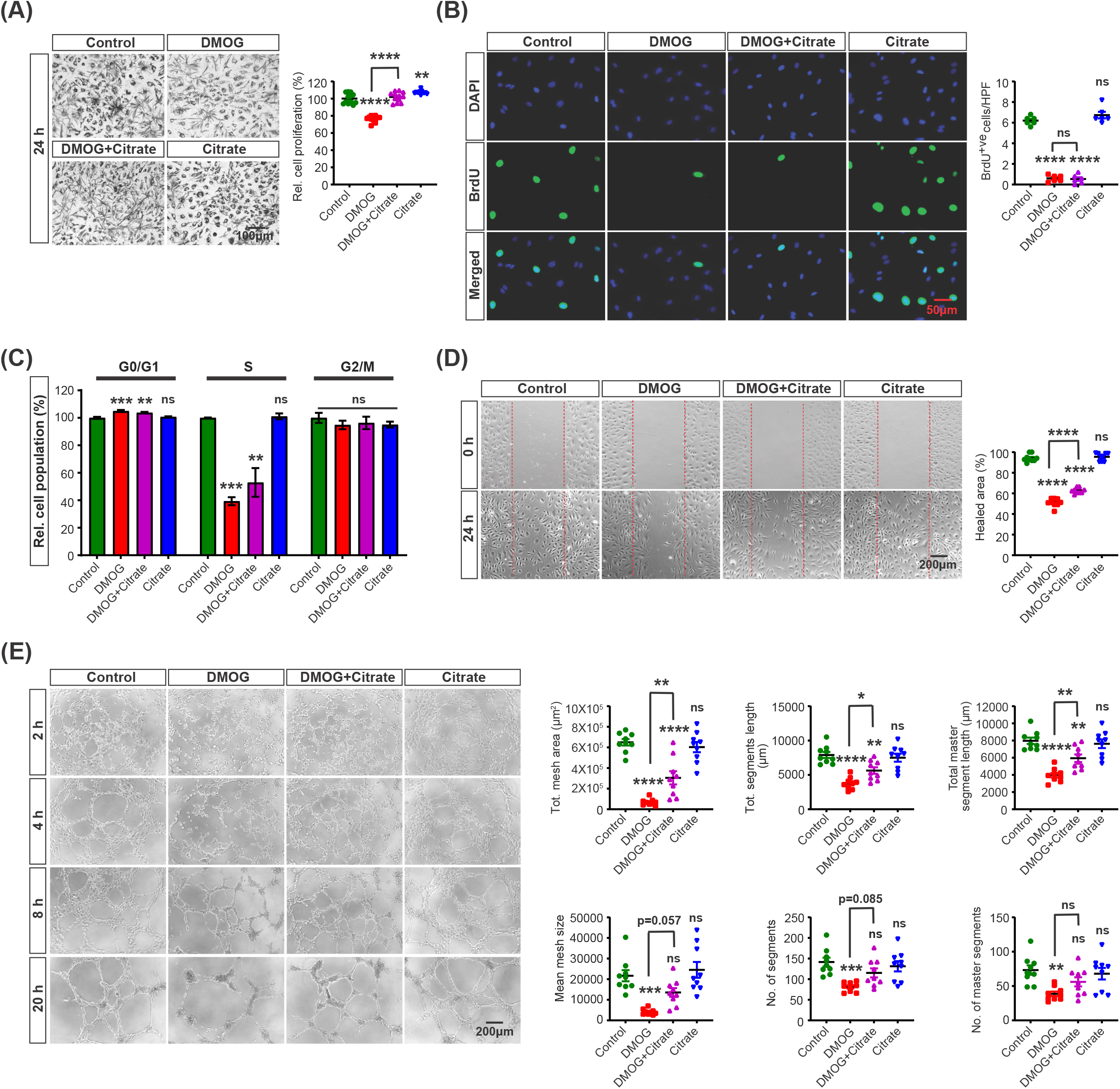
Citrate supplementation partially restores the DMOG-induced defects in endothelial migration and tube formation capacity. (**A**) Representative bright field images of formazan crystal formed after MTT incubation with control, DMOG (1mM), DMOG + citrate and citrate (0.5mM) treated HPAEC. Right graph shows relative HPAEC proliferation calculated by MTT assay. (**B**) Representative images of BrdU immunostaining under the conditions indicated in A. Right graph shows semi-quantitative analysis of BrdU positive cells per hpf. (**C**) Quantitative analysis of cell cycle showing relative percentage of cell population in G0/G1, S and G2/M cell cycle phase. (**D**) Representative images of 2D scratch wound assay of control or DMOG-treated cells and semi-quantitative analysis of healed area after 24 h. (**E**) Representative images of tubes formed at the indicated time points in control, DMOG, DMOG + citrate and citrate treated cells and semi-quantitative analysis of different parameters at 20 h time point. Data are pooled from 3 independent experiments and represented as mean ± SEM. Statistics were determined by one-way ANOVA with Sidak correction for multiple comparisons. *, P<0.05; **, P<0.01; ***, P<0.001; ****, P<0.0001; ns, not significant. Asterisks above bars indicate significant difference between control and treated group, whereas asterisks above lines indicate significant difference between DMOG and DMOG + citrate treated groups.

**Figure 5.**
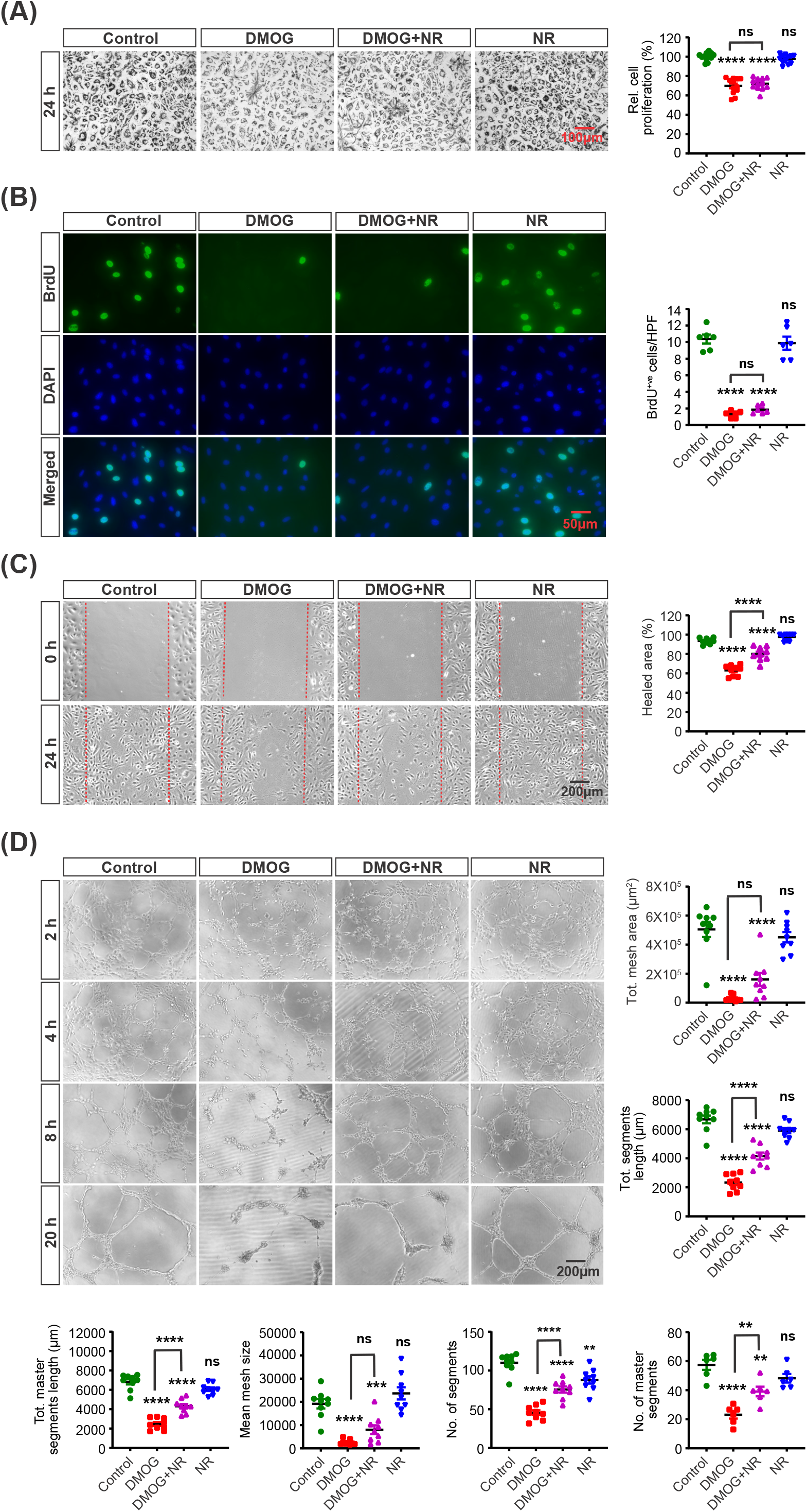
Nicotinamide riboside supplementation partially rescues the DMOG-induced defects in endothelial migration and tube formation capacity. (**A**) Representative bright field images of formazan crystal formed after incubation with MTT in control, DMOG (1 mM), DMOG + NR and NR (200 µM) treated HPAEC. Right graph shows relative HPAEC proliferation assessed by MTT assay. (**B**) Representative images of BrdU immunostaining. Right graph shows semi-quantitative analysis of BrdU positive cells/hpf. (**C**) Representative images of 2D scratch wound assay and semi-quantitative analysis of healed area after 24 h. (**D**) Representative images of tubes formed at different time points in control, DMOG, DMOG + NR and NR treated ECs and semi-quantitative analysis of different parameters at 20 h time point. Data are pooled from 3 independent experiments and represented as mean ± SEM. Statistics were determined by one-way ANOVA with Sidak correction for multiple comparisons. **, P<0.01; ***, P<0.001; ****, P< 0.0001; ns, not significant. Asterisks above bars indicate significant difference between control and treated group, whereas asterisks above lines indicate significant difference between DMOG and DMOG + NR treated groups. NR, nicotinamide riboside.

## METHODS

### Cell culture

Human primary pulmonary artery endothelial cells (HPAEC) were purchased from ATCC (ATCC) and maintained in endothelial cell growth medium-2 (EGM-2, Lonza) supplemented with EGM-2 SingleQuot Kit (Lonza). HPAEC were cultured in multi-well cell culture plates and flasks coated with 0.1 % gelatin. All experiments were performed in cells cultured for ≤7 passages. DMOG (Tocris) was dissolved in dimethyl sulfoxide (DMSO). Trimethyl citrate (Sigma), methyl pyruvate (Thermo Scientific™), methyl aspartate (Santa Cruz) and nicotinamide riboside (Cayman) were used for metabolic repletion experiments.

### Proliferation assays

Cell proliferation was assessed by 3-(4,5-dimethylthiazol-2-yl)-2,5-diphenyltetrazolium bromide (MTT) assay as described by Zirkel A et al. (35) and BrdU incorporation assay. Briefly, HPAEC were seeded in gelatin-coated 96-well culture plates. The next day, cells were treated with DMOG (1 mM), while control cells were treated with DMSO. After 20 h incubation, MTT solution (Invitrogen) was added for 4 h at 37°C. Subsequently, the intracellular formazan crystals were solubilized with DMSO and the absorbance of the colored solution was read at 540 nm. For BrdU labeling, cells were grown in 4-well chamber slides. Control and DMOG-treated cell were incubated with BrdU solution (BD Pharmingen™) for 3 h under optimal growth conditions. Post incubation, cells were washed 3 times with 0.1% tween 20 in PBS (PBST) and fixed with a methanol/acetone mixture (molar ratio 1:1) for 30 minutes. After a wash, cells were incubated with 1.5 M HCl for 25 minutes at room temperature. Next, cells were washed again with PBST and blocked with 3% BSA for 1 h. After blocking, cells were incubated with an anti-BrdU primary antibody (Cell Signaling Technology) overnight at 4 °C. After 3 washes with PBST, cells were probed with a secondary antibody (Abcam) at 37°C for 1 h. Finally, cells were washed and mounted with Vectashield Vibrance Antifade Mounting Medium containing DAPI (Vector Laboratories, Inc.). Five random images per well were captured using a fluorescence microscope (Leica DM IL LED) and BrdU positive nuclei were counted. Minimum of 3 independent experiments were performed and analyzed.

### 2D scratch migration assay

Cell migration was assessed using a scratch wound healing assay (36). HPAEC were seeded in 12-well plates and allowed to grow until monolayer formation. Scratches were created manually using pipette tip, washed with media followed by addition of DMOG or DMSO. To assess cell migration, bright field images were captured by at 0 h and 24 h time points. Percentage of healed area was measured using Fiji software based on ImageJ (NIH).

### Matrigel tube formation assay

Tube formation assay was performed using the angiogenesis assay kit (Abcam). Briefly, HPAEC treated with DMOG or DMSO for 24 hours were trypsinized and resuspended in EGM-2 media containing either DMOG or DMSO and then added to Matrigel coated 96-well plates. Bright field images were captured at different time points (2, 4, 8, 20 h). Captured images were analyzed for total meshes area, mean mesh size, branching intervals, number of segments, number of master segments, total segment length, and total master segment length using the image J-based Angiogenesis Analyzer tool.

### Cell cycle analysis and Annexin V/ PI cell death assays

Following 24 hours treatment with DMOG, cells were trypsinized, centrifuged at 300g for 5 minutes. Cell pellets were resuspended in ice cold PBS, centrifuged and fixed overnight with 70% ethanol. Fixed cells were resuspended in propidium iodide (PI) staining buffer (Invitrogen) and incubated at 37°C for 30 min. Subsequently, cell cycle was assessed by a BD LSR Fortessa flow cytometer (BD Biosciences) using FACS Diva and FlowJo software. Cell death was examined by a FITC Annexin V Apoptosis Detection Kit I (BD Pharmingen™) following manufacturer’s protocol. ECs were trypsinized, centrifuged, and washed twice with cold PBS and resuspended in a Binding Buffer provided with the kit. For flow cytometric preparation, cell suspensions were incubated with 5 µL of Annexin V-FITC and 5 μL of PI for 15 minutes at room temperature in the dark. After incubation, 400 μL of Binding Buffer was added and applied to the BD LSR Fortessa flow cytometer (BD Biosciences). Data analysis was performed using FACS Diva and FlowJo software.

### Isolation of nuclear protein and immunoblotting

Nuclear protein was isolated using NE-PER Nuclear and Cytoplasmic Extraction Reagents (Thermo Fisher Scientific) following manufacturer’s instructions. Nuclear protein extracts were separated on SDS-PAGE gel, transferred on nitrocellulose membrane and incubated with HIF-1α (Cayman) or HIF-2α (Novus Biologicals) antibodies at 4°C. After overnight incubation, nitrocellulose membrane was washed and incubated with secondary antibody (Novus Biologicals) for 90 minutes at 4°C. Blots were visualized by SuperSignal™ West Femto Chemiluminescent Substrate (Thermo Fisher Scientific) and chemiluminescence was detected on an iBright Western Blot Imaging Systems (Thermo Fisher Scientific).

### HIF-1_α_ and HIF-2_α_ Immunostaining

Cells were fixed with Image-iT™ Fixative Solution (Thermo Fisher Scientific) for 10 minutes at room temperature. Subsequently, cells were washed 3 times with PBS, permeabilized for 10 minutes with 0.1 % tween 20 in PBS and then nonspecific antigens were blocked with 3% BSA for 2 h. After blocking, cells were incubated with primary antibody against HIF-1α (Cayman) or HIF-2α (Novus Biologicals) at 4 °C overnight, followed by incubation with secondary antibody (Thermo Fisher Scientific) for 2 h at room temperature. Following 3 washes with PBS, cells were mounted with Vectashield Vibrance Antifade Mounting Medium containing DAPI (Vector Laboratories, Inc.). Images were captured by a fluorescence microscope (Leica DM IL LED) and analyzed using Fiji software.

### RNA isolation and RNA-sequencing

Total RNA was extracted using RNeasy Mini Kit (Qiagen) following the manufacturer’s instructions. RNA integrity was checked with Agilent Technologies 2100 Bioanalyzer. Poly(A) tail-containing mRNAs were purified using oligo-(dT) magnetic beads with two rounds of purification. After purification, poly(A) RNA was fragmented using divalent cation buffer in elevated temperature. Poly(A) RNA sequencing library was prepared following Illumina’s TruSeq-stranded-mRNA sample preparation protocol. Quality control analysis and quantification of the sequencing library were performed using Agilent Technologies 2100 Bioanalyzer High Sensitivity DNA Chip. Paired-ended sequencing was performed on Illumina’s NovaSeq 6000 sequencing system. Adapter contaminated peaks, low quality bases and undermined bases were removed using Cutadapt (37) and perl scripts. Sequence quality was verified using FastQC (http://www.bioinformatics.babraham.ac.uk/projects/fastqc/). HISAT2 (38) was used to map reads to the genome of ftp://ftp.ensembl.org/pub/release-96/fasta/homo_sapiens/dna/ and mapped reads of each sample were assembled using StringTie (39). Subsequently, all transcriptomes were merged to reconstruct a comprehensive transcriptome using perl scripts and gffcompare. Final transcriptome was analyzed using StringTie (39) and edgeR (40) to estimate the expression level of all transcripts. StringTie was used to perform expression level for mRNAs by calculating FPKM. The differentially expressed protein coding mRNAs were selected with fold change >1.5 and with adjusted P value < 0.05 by R package edgeR. Volcano plot was generated by EnhancedVolcano (https://github.com/kevinblighe/EnhancedVolcano). Enrichment analyses was performed in EnrichR (https://maayanlab.cloud/Enrichr/) and pathfindR (41).

### Metabolomic profiling

To assess the effect of DMOG on HPAEC, global metabolic profiles were determined in cell pellets and supernatants derived from HPAEC grown in the presence of DMOG or vehicle. The metabolomic analysis was conducted by Metabolon, Inc (Durham, NC). Samples were prepared using the automated MicroLab STAR® system (Hamilton Company, USA). A recovery standard was added prior to the first step in the extraction process for QC purposes. For the maximum recovery of diverse metabolites, proteins were precipitated with methanol under vigorous shaking for 2 min (Glen Mills GenoGrinder 2000) followed by centrifugation. The resulting extract was divided into five fractions: one for analysis by UPLC-MS/MS with positive ion mode electrospray ionization, one for analysis by UPLC-MS/MS with negative ion mode electrospray ionization, one for analysis by UPLC-MS/MS polar platform (negative ionization), one for analysis by GC-MS, and one sample was reserved for backup. Samples were placed briefly on a TurboVap® (Zymark) to remove the organic solvent. For LC, the samples were stored overnight under nitrogen before preparation for analysis. For GC, each sample was dried under vacuum overnight before preparation for analysis. Several types of controls were analyzed in concert with the experimental samples: a pooled matrix sample generated by taking a small volume of each experimental sample served as a technical replicate throughout the data set; extracted water samples served as process blanks; and a cocktail of QC standards that were carefully chosen not to interfere with the measurement of endogenous compounds were spiked into every analyzed sample, allowing instrument performance monitoring and aiding chromatographic alignment. Instrument variability was determined by calculating the median relative standard deviation (RSD) for the standards that were added to each sample prior to injection into the mass spectrometers. Overall process variability was determined by calculating the median RSD for all endogenous metabolites (i.e., non-instrument standards) present in 100% of the pooled matrix samples.

Four major components were involved in the informatics system: Laboratory Information Management System (LIMS), the data extraction and peak-identification software, data processing tools for QC and compound identification, and a collection of information interpretation and visualization tools. LAN backbone, and a database server running Oracle 10.2.0.1 Enterprise Edition worked as the hardware and software foundation for the components of the informatics system.

Raw data was extracted, peak-identified and QC processed using Metabolon’s hardware and software. Compounds were identified by comparison to library entries of purified standards or recurrent unknown entities based on authenticated standards that contains the retention time/index (RI), mass to charge ratio (*m/z)*, and chromatographic data (including MS/MS spectral data) on all molecules present in the library. Furthermore, biochemical identifications are based on three criteria: retention index within a narrow RI window of the proposed identification, accurate mass match to the library +/- 0.005 amu, and the MS/MS forward and reverse scores between the experimental data and authentic standards. For quantification, peaks were quantified using area-under-the-curve and were normalized. Significantly altered metabolites (P<0.05) were used to perform metabolite sets enrichment analysis by MetaboAnalyst (https://www.metaboanalyst.ca).

### Extraction and measurement of NAD^+^

Cells were grown in 10 cm^2^ dishes. Following 24 h of treatment with DMOG or vehicle, media was aspirated, and cells were quickly rinsed with ice cold normal saline. Culture dishes were placed on dry ice and 600µL ice cold extraction buffer (40:40:20 acetonitrile/methanol/200 mM NaCl, 10 mM Tris-HCl, pH 9.2) were added directly to the cells. Cells were scraped and transferred to 1.5 mL tubes and were frozen in liquid nitrogen. Cells were subjected to two freeze thaw cycles with vortex of 15 seconds after each thaw. After lysis, cell lysates were centrifuged at 20,000xg for 15 minutes at 4°C. Metabolite containing supernatant was transferred to fresh 1.5 mL tube and protein pellets were used for protein quantification. The metabolite containing supernatants were dried down by a SpeedVac concentrator (Thermo Savant). After reconstitution in 60% acetonitrile, samples were analyzed by Thermo Fisher Scientific U3000 UHPLC equipped with Q Exactive™ Q-Orbitrap MS.

## QUANTIFICATION AND STATISTICAL ANALYSIS

Statistical analyses were performed using the program “R” (http://cran.r-project.org/) or GraphPad Prism 9 (GraphPad Software, La Jolla, CA). Two-tailed t-test with Welch’s correction was used to calculate statistical significance between two groups, while for multiple comparisons statistical significance was calculated by one-way ANOVA with Sidak correction. A two-sided significance level of 5% was considered statistically significant. RNA-seq data was analyzed using the software R with package edgeR. Adjusted P values < 0.05 were considered statistically significant. All the experiments were performed at least 3 times independently. Results are reported as mean ± SEM.

## DISCUSSION

Here, we discovered a previously unidentified role for PHD inhibition in suppressing angiogenic properties of human vascular endothelium, which may impair vascular homeostasis and regeneration in ischemic tissues. Our studies demonstrate that induction of EC pseudohypoxia by PHD inhibition impairs EC proliferation, migration, and tube formation capacities. Although metabolic reprogramming induced by hypoxic signaling has been implicated in vascular biology (12, 42), the role of PHD/HIF pathway in endothelial metabolism remains unclear. By combining unbiased RNA-seq and metabolomic analysis, we found that PHD inhibition induces an unfavorable metabolic reprogramming for angiogenesis characterized by alterations in amino acid metabolism and suppression of mitochondrial metabolism. Furthermore, our mechanistic studies revealed that suppression of mitochondrial metabolism with reduced NAD^+^ regeneration may at least partially drive the defects in migration and tube forming capacity of EC in the context of PHD inhibition. These findings demonstrate a maladaptive metabolic reprogramming as important molecular signature that characterize the response of vascular endothelium to chemical PHD inhibition.

In hypoxic tissues, parenchymal cells respond to oxygen deprivation via HIFs, which up-regulate angiogenic genes leading to activation of ECs and vessel growth (12, 43, 44). As oxygen sensors, PHDs have been also implicated in regulating angiogenesis mostly through their role in controlling the stability of HIFs (45). Using DMOG, a 2-oxoglutarate analogue which competitively inhibits PHDs, we show that chemical PHD inhibition suppresses angiogenesis by reducing EC proliferation, migration and tube formation. These findings were quite surprising, particularly since prior studies have shown that both HIF-1 and HIF-2 in EC promote angiogenesis in a cell autonomous manner (23, 46). Notably, germline deletion of *Phd2* results in abnormal placental vascularization and embryonic lethality (47), while heterozygosity for *Phd2* does not alter capillary density despite the increase in HIF activity (48). Furthermore, our findings agree with prior study, which showed that exposure to hypoxia slowed EC growth, decreases entry of cell into S phase and delayed wound repair (49). While our study did not find an increase in EC death by DMOG, other studies have shown that hypoxia may trigger EC apoptosis by induction of NFkB (50) or HIF-1 dependent stabilization of p53 (51). On the other hand, several studies have reported that moderate hypoxia promotes EC proliferation, survival, and migration (12). Therefore, inhibition of PHDs may differentially regulate angiogenesis depending on the context, means of inhibition and cell-type affected.

Herein, we explored the transcriptional responses of EC to HIF-1 and HIF-2 stabilization with the prolyl-hydroxylase inhibitor DMOG. Compared to prior microarray-based studies, we employed RNA-seq as an unbiased approach to uncover a wide range of transcriptional responses that may drive EC functional alterations to chemical PHD inhibition. As expected, we found that DMOG exposure mimics hypoxia and regulates the expression of several angiogenic genes. Related to decreased EC proliferation following DMOG exposure, our RNA-seq analysis showed reduced transcription of genes associated with MCM complex formation, which is necessary for DNA synthesis and cell proliferation (24). This finding is consistent with the study by Hubbi, which reported that HIF-1α inhibits MCM protein complex activation leading to decreased DNA replication and cell cycle arrest (52). Consistently, we found suppression of transcripts encoding DNA polymerases *DNA polymerase alpha 1, catalytic subunit* (*POLA1*), *DNA polymerase alpha 2, accessory subunit (POLA2*), *DNA polymerase epsilon 2, accessory subunit (POLE2*), *DNA polymerase delta 3, accessory subunit* (*POLD3*), an effect that may contribute to the antiproliferative action of DMOG on endothelial cells.

Endothelial metabolism has emerged as a critical regulator of EC homeostasis and growth (45). While EC generate most of their ATP by glycolysis, PHD inhibition led to a broad metabolic reprogramming as demonstrated by our transcriptomic and metabolomic analysis. Besides the expected upregulation of glycolytic genes by DMOG, we found significant transcriptional suppression for several subunits of MC1 complex and its product NAD^+^. Previously, hypoxia has been reported to inhibit MC1 in different cell types including HUVECs (53) by induction of *NADH dehydrogenase [ubiquinone] 1 alpha subcomplex, 4-like 2* (*NDUFA4L2*). Nevertheless, we found no up-regulation of *NDUFA4L2* by DMOG, which suggests a distinct mechanism for suppression of MC1. Remarkably, the suppression of MC1 could drive the low NAD^+^/NADH ratio, which impairs flow of TCA cycle and glycolytic flux (25). Furthermore, we found increase in NAD^+^ levels following repletion with citrate, that may be supported by cytosol confined NADH recycling through malate dehydrogenase 1 (MDH1), a process reported to operate in cells with mitochondrial dysfunction (34). Finally, the observed improvement of EC migration and tube formation by NR or citrate supplementation is probably due to NAD^+^ -induced enhancement of glycolytic flux, as glycolysis and not oxidative phosphorylation is the main regulator for EC migration (2, 6).

Consistent with our findings is a recent study in which S-2-hydroxyglutarate (S-2HG), a metabolite with inhibitory effect on PHDs, lowered the proliferative capacity of EC, an effect also recapitulated by genetic PHD inactivation (54). The precise metabolic mechanism by which chemical PHD inactivation impairs proliferation of human ECs remains to be defined but probably involves inability to generate the essential metabolites for macromolecule synthesis due to suppressed mitochondrial oxidation and glutamine metabolism. While reduced aspartate synthesis is critical for cell proliferation upon electron transport chain (ETC) inhibition (26, 27), its supplementation, like that of pyruvate, failed to improve the proliferative defect of DMOG-treated ECs. Furthermore, the inhibition of glutamine metabolism by DMOG is likely to play a major role in the reduced proliferative capacity of DMOG-treated cells, as glutamine supplies most carbons in TCA cycle and contributes to biomass synthesis in ECs (55). Interestingly, a recent study showed a HIF-independent mechanism by which DMOG impairs glutamine metabolism, which involves the inhibition of multiple enzymatic steps, including glutamate dehydrogenase (GDH) by inducing the generation of the metabolite *N*-oxalylglycine (NOG) (56).

Recent advances in our understanding of endothelial metabolism in health and disease have created an expectation that modulation of EC metabolism by hypoxia signaling-based therapeutics may have promising clinical potential. Nevertheless, our study shows that chemical PHD inhibition by DMOG induces an unfavorable metabolic reprogramming for angiogenic competence. Future studies should investigate the role of PHDs in the endothelium and define HIF-dependent vs independent effects on EC metabolism.

### Limitations of Study

This study has several limitations. First, like other studies, we used DMOG to inactivate PHDs and activate HIF signaling, but DMOG may have unpredicted off-target effects by inhibiting other members of 2-oxoglutarate-dependent dioxygenases. Future studies should explore the effects of other chemical PHD inhibitors and compare them to genetic PHD inactivation. Second, we did not dissect the role of HIF-1α and HIF-2α on the DMOG-induced defects on angiogenic competence and EC metabolism. Finally, more studies should be performed to investigate the role of endothelial PHDs on angiogenic responses in vivo as the tissue specific microenvironment would be critical in this regard.

## AUTHOR CONTRIBUTIONS

PPK and RT conceived the study; RT, NC and PPK designed the experiments; RT and PVB performed experiments; RT, PVB, PG, MJS, YZ, SEQ, NC and PPK analyzed and interpreted data; PG developed and optimized the quantitative LC/MS assay for measurement of NAD^+^; RT and PPK wrote the manuscript and all authors approved and commented on the manuscript.

## ACKNOWLEDGMENTS

This work was supported by National Institutes of Health (NIH) grant R01DK115850 (PPK) and the Kansas INBRE, P20 GM103418 (RT). Metabolomics services were performed by the Metabolomics Core Facility at Robert H. Lurie Comprehensive Cancer Center of Northwestern University. We acknowledge the Northwestern University George M. O’Brien Kidney Research Core Center (NU GoKidney), an NIH/NIDDK funded program (P30 DK114857) for their core services and support. The funders had no role in study design, data collection and interpretation, or the decision to submit the work for publication.

## TITLE FOR EXCEL FILE

**Table S1, Related to Figure 3. Metabolomics analysis of cell lysates and media of HPAEC treated with DMOG compared to vehicle**. Shown are fold changes with corresponding *p*-and *q*-values (5 DMOG-treated and 5 vehicle-treated samples). Statistically significant increased metabolites are marked in red while green is used for significantly decreased metabolites. Data were collected using the Metabolon LC/MS and GC/MS platforms.

