## Supplemental Figures for "Chemical inhibition of prolyl hydroxylases impairs angiogenic competence of human vascular endothelium through metabolic reprogramming"

**Supplementary Figure 1. DMOG treated endothelial cells maintain viability and stabilize HIF-α (related to Figure 1).** (**A**) Representative images of immunostaining of HIF-1α and HIF-2α proteins in control and DMOG-treated cells. (**B**) Immunoblot analysis of HIF-1α and HIF-2α in nuclear extracts isolated from control and DMOG-treated cells. (**C**) Shown are percentages of Annexin V/propidium iodide (PI) positive cells after 24 h treatment with vehicle (control) or DMOG (1 mM). Data are pooled from 3 independent experiments and represented as mean ± SEM. Statistics were determined by two-tailed t-test. **, $P<0.01$.

### Supplemental Figure 1

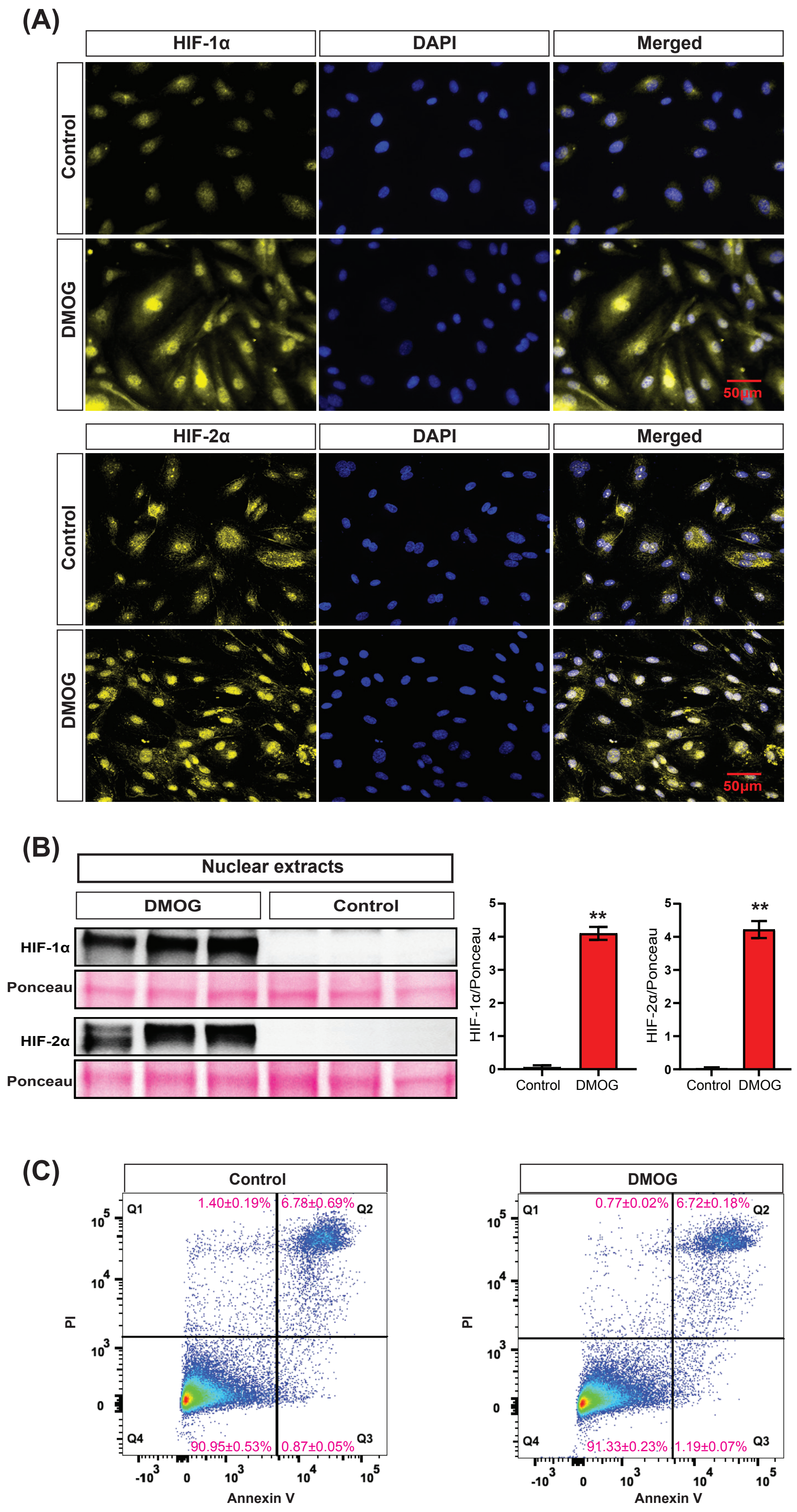

**Supplementary Figure 2. Prolyl hydroxylase inhibition by DMOG induces broad transcriptional reprogramming in endothelial cells (related to Figure 2).** (**A**) Volcano plot of RNA-seq data showing number of significantly up-regulated and down-regulated genes in DMOG-treated cells compared to control. (**B**) Heatmaps showing the top 50 up-regulated (top) and top 50 down-regulated genes (bottom) in the transcriptome of HPAECs treated with DMOG for 24h. (**C**) Pathway map illustrating transcriptional changes induced by DMOG in genes involved in glycolysis, TCA cycle and ETC. Up-regulated and down-regulated genes by DMOG are shown in red and blue color respectively, whereas genes in black color showed no significant transcriptional changes. TCA, tricarboxylic acid cycle; ETC, electron transport chain.

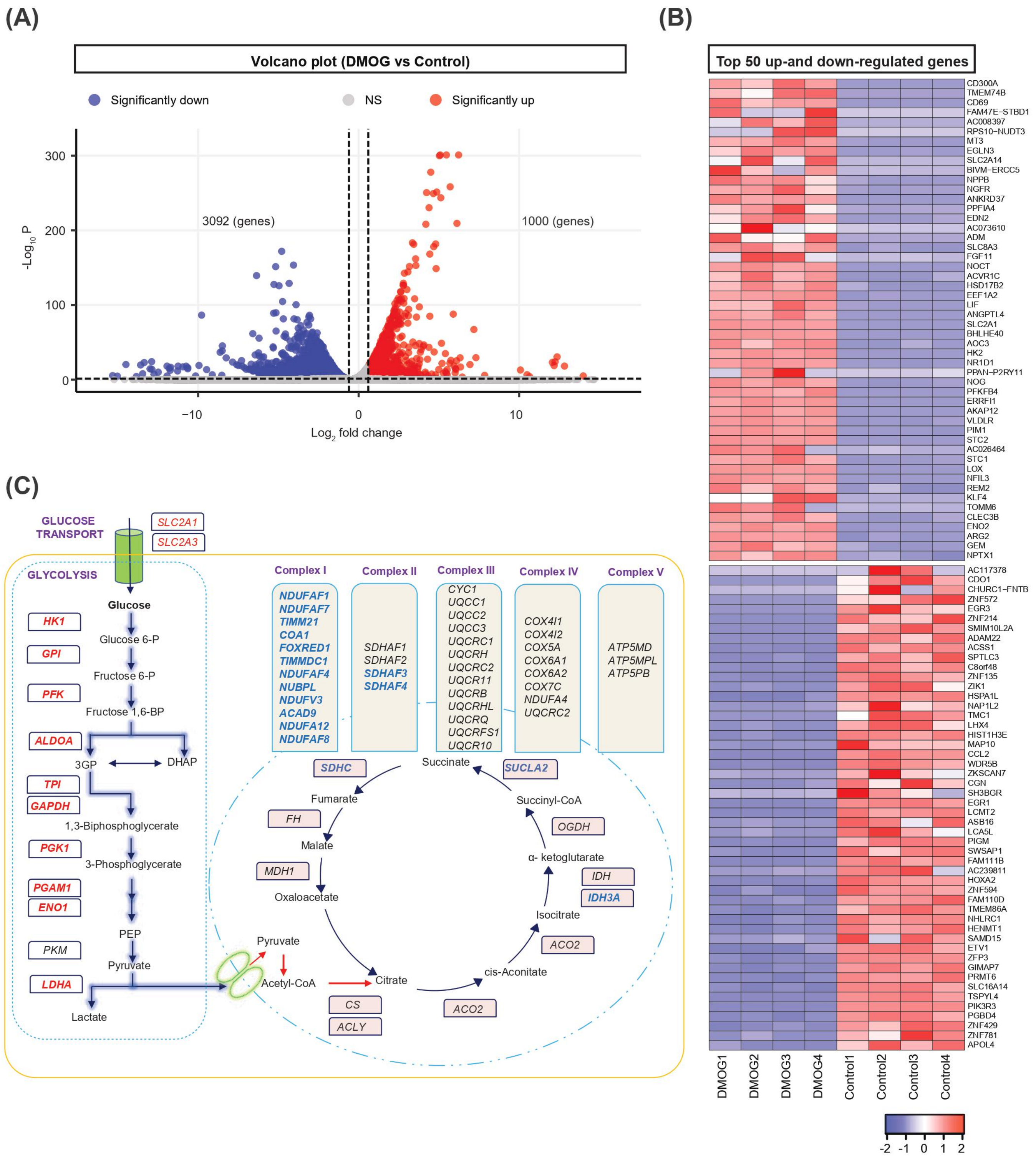
(A)
Volcano plot (DMOG vs Control)
Significantly down
NS
Significantly up
3092 (genes)
1000 (genes)
-Log10 P
Log2 fold change
300
200
100
0
−10
0
10
(B)
Top 50 up-and down-regulated genes
CD300A
TMEM74B
CD69
FAM47E−STBD1
AC008397
RPS10−NUDT3
MT3
EGLN3
SLC2A14
BIVM−ERCC5
NPPB
NGFR
ANKRD37
PPFIA4
EDN2
AC073610
ADM
SLC8A3
FGF11
NOCT
ACVR1C
HSD17B2
EEF1A2
LIF
ANGPTL4
SLC2A1
BHLHE40
AOC3
HK2
NR1D1
PPAN−P2RY11
NOG
PFKFB4
ERRFI1
AKAP12
VLDLR
PIM1
STC2
AC026464
STC1
LOX
NFIL3
REM2
KLF4
TOMM6
CLEC3B
ENO2
ARG2
GEM
NPTX1
AC117378
CDO1
CHURC1−FNTB
ZNF572
EGR3
ZNF214
SMIM10L2A
ADAM22
ACSS1
SPTLC3
C8orf48
ZNF135
ZIK1
HSPA1L
NAP1L2
TMC1
LHX4
HIST1H3E
MAP10
CCL2
WDR5B
ZKSCAN7
CGN
SH3BGR
EGR1
LCMT2
ASB16
LCA5L
PIGM
SWSAP1
FAM111B
AC239811
HOXA2
ZNF594
FAM110D
TMEM86A
NHLRC1
HENMT1
SAMD15
ETV1
ZFP3
GIMAP7
PRMT6
SLC16A14
TSPYL4
PIK3R3
PGBD4
ZNF429
ZNF781
APOL4
DMOG1
DMOG2
DMOG3
DMOG4
Control1
Control2
Control3
Control4
-2 -1 0 1 2
(C)
GLUCOSE TRANSPORT
SLC2A1
SLC2A3
GLYCOLYSIS
Glucose
HK1
Glucose 6-P
GPI
Fructose 6-P
PFK
Fructose 1,6-BP
ALDOA
3GP
DHAP
TPI
GAPDH
1,3-Biphosphoglycerate
PGK1
3-Phosphoglycerate
PGAM1
ENO1
PEP
PKM
Pyruvate
LDHA
Lactate
Complex I
NDUFAF1
NDUFAF7
TIMM21
COA1
FOXRED1
TIMMDC1
NDUFAF4
NUBPL
NDUFV3
ACAD9
NDUFA12
NDUFAF8
Complex II
SDHAF1
SDHAF2
SDHAF3
SDHAF4
Complex III
CYC1
UQCC1
UQCC2
UQCC3
UQCRC1
UQCRH
UQCRC2
UQCR11
UQCRB
UQCRHL
UQCRQ
UQCRFS1
UQCR10
Complex IV
COX4I1
COX4I2
COX5A
COX6A1
COX6A2
COX7C
NDUFA4
UQCRC2
Complex V
ATP5MD
ATP5MPL
ATP5PB
SDHC
Succinate
SUCLA2
Fumarate
Succinyl-CoA
FH
OGDH
Malate
α- ketoglutarate
MDH1
IDH
IDH3A
Oxaloacetate
Isocitrate
Pyruvate
ACO2
Acetyl-CoA
Citrate
cis-Aconitate
CS
ACLY
ACO2

**Supplementary Figure 3**. **Metabolomic analysis of DMOG treated ECs compared to control (related to Figure 3).** (**A**) Experimental workflow for the metabolomic analysis of cellular lysates and media from control and DMOG treated HPAEC. Shown in (**B**) and (**C**) are biochemical importance plots generated by random forest classification of metabolites measured in cells and media, respectively from control and DMOG-treated HPAEC.

#### Supplemental Figure 3

**(A)**

| Groups | Matrix (n) Cells | Matrix (n) Media |
|---|---|---|
| Control | 5 | 5 |
| DMOG | 5 | 5 |

UPLC-MS/MS | GC/MS

| Welch's Two-sample t-test | DMOG/Control Cells | DMOG/Control Media |
|---|---|---|
| Total biochemicals (p ≤ 0.05) | 126 | 45 |
| Biochemicals (↑↓) | 70 \| 56 | 19 \| 26 |

**(B)**

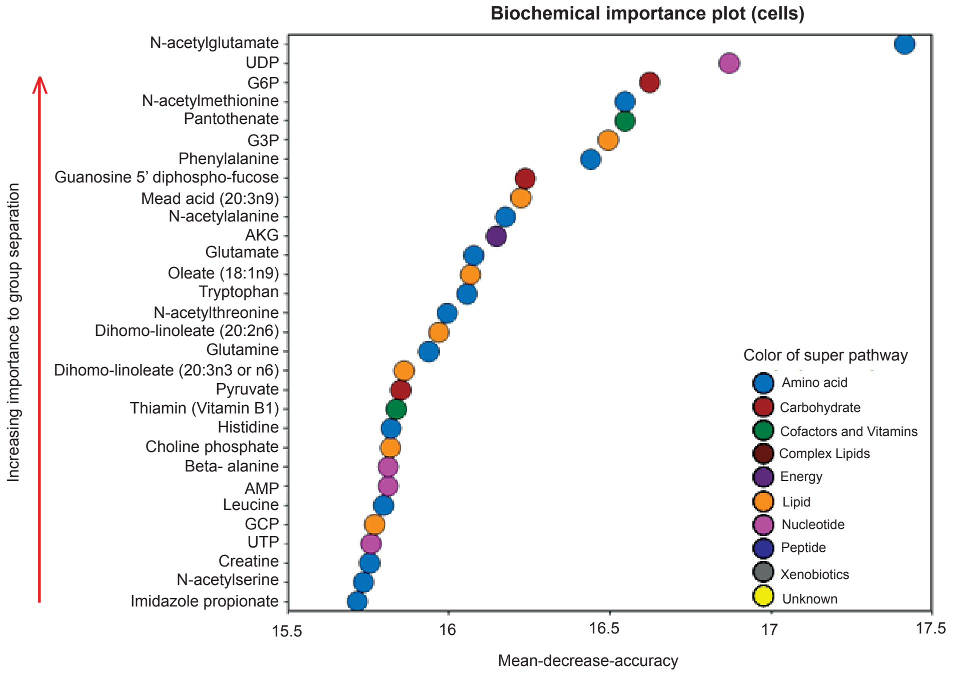


**(C)**

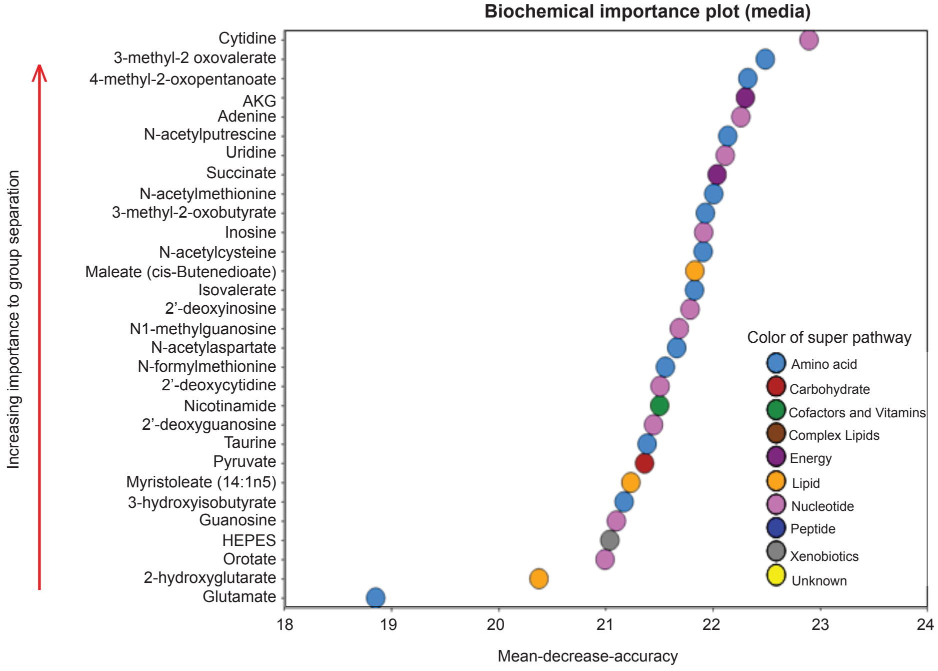

**Supplementary Figure 4. Supplementation of membrane permeable derivatives of pyruvate (methyl pyruvate) and aspartate (methyl aspartate) do not affect angiogenic defects induced by DMOG (related to Figure 4).** (**A**) Effect of pyruvate supplementation (2 mM, 1 mM and 0.5 mM) on HPAEC proliferation assessed by MTT assay. (**B**) Representative images of 2D scratch wound assay of control, DMOG, DMOG + pyruvate and pyruvate (0.5mM) treated cells and semi-quantitative analysis of healed area after 24h. (**C**) Effect of aspartate supplementation (5mM, 2mM and 0.5mM) on HPAEC proliferation assessed by MTT assay. (**D**) Representative images of 2D scratch wound assay of control, DMOG, DMOG + aspartate and aspartate (0.5mM) treated cells and semi-quantitative analysis of healed area after 24 h. (**E**) Representative images of tubes formed at indicated time points in control, DMOG, DMOG + pyruvate, DMOG + aspartate, pyruvate (0.5 mM) and aspartate (0.5 mM) treated cells and semi-quantitative analysis of different parameters at 20 h time point. Data are pooled from 3 independent experiments and represented as mean ± SEM. Statistics were determined by one-way ANOVA with Sidak correction for multiple comparisons. ***, $P<0.001$; ****, $P< 0.0001$; ns, not significant. Asterisks/ns above bars indicate significance level between control and treated group, whereas asterisks/ns above the lines indicate significance level between DMOG and DMOG + pyruvate or DMOG + aspartate treated groups.

#### Supplemental Figure 4

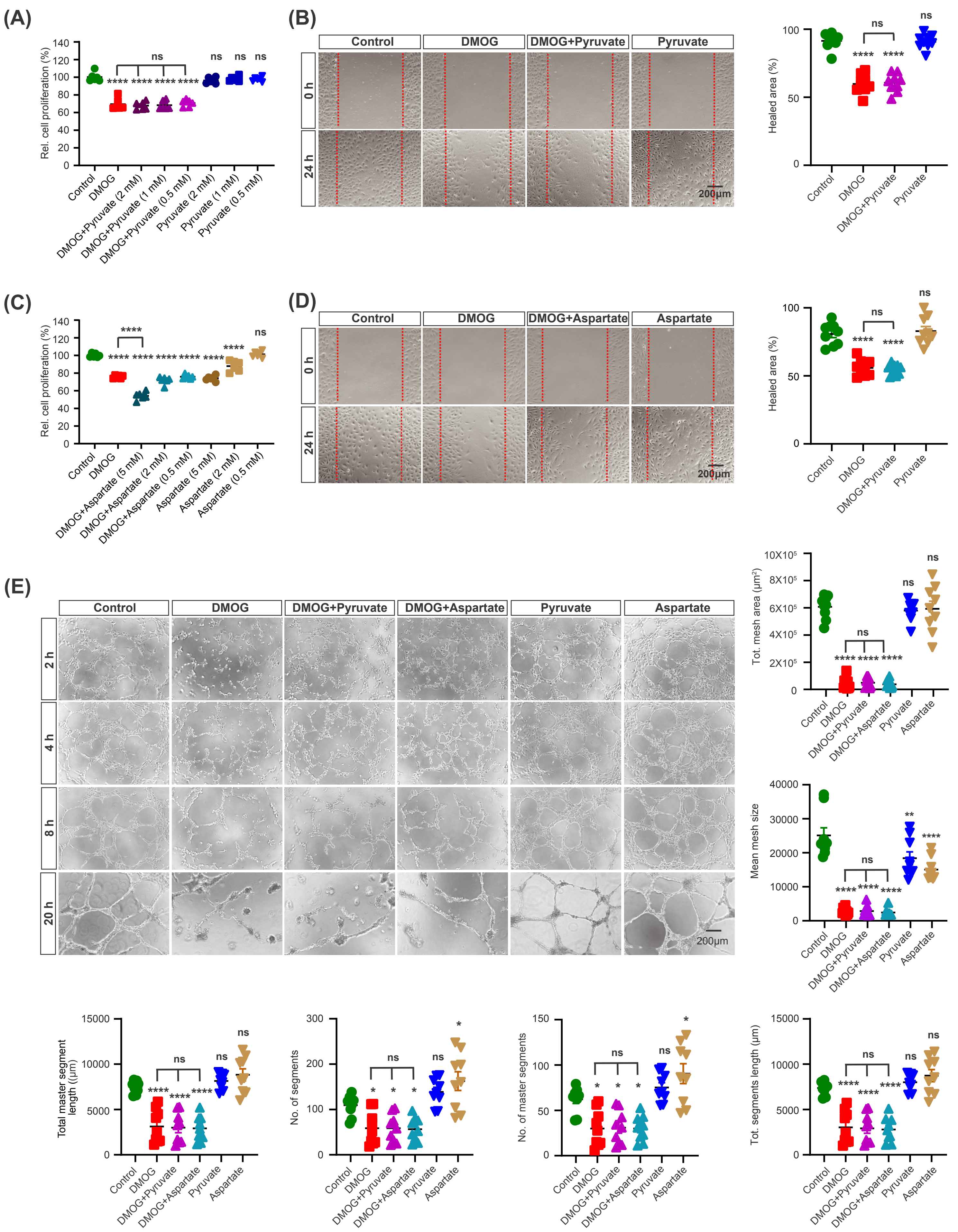

**Supplementary Figure 5. Effect of citrate on cell cycle progression, HIF-α protein and $NAD^+$ levels (related to Figure 4).** HPAEC were treated with different compounds for 24 h. (**A**) Histogram of cell cycle analysis for control, DMOG, DMOG + citrate, and citrate (0.5 mM) treated cells. (**B**) Representative images of immunostaining for HIF-1α and HIF-2α proteins in control, DMOG, DMOG + citrate and citrate treated cells. (**C**) Immunoblot analysis of HIF-1α and HIF-2α in nuclear extracts isolated from control, DMOG, DMOG + citrate, and citrate treated cells. (**D**) Relative abundance of $NAD^+$ levels in cellular extracts from control, DMOG, DMOG + citrate, and citrate treated cells. Data are represented as mean ± SEM. Statistics were determined by one-way ANOVA with Sidak correction for multiple comparisons. *, P<0.05; **, P<0.01; ***, P<0.001; ****, P< 0.0001; ns, not significant. Asterisks/ns above bars indicate significance level between control and treated group, whereas asterisks/ns above lines indicate significance level between DMOG and DMOG + citrate treated groups. D + Citr., DMOG + citrate treated group.

#### Supplemental Figure 5

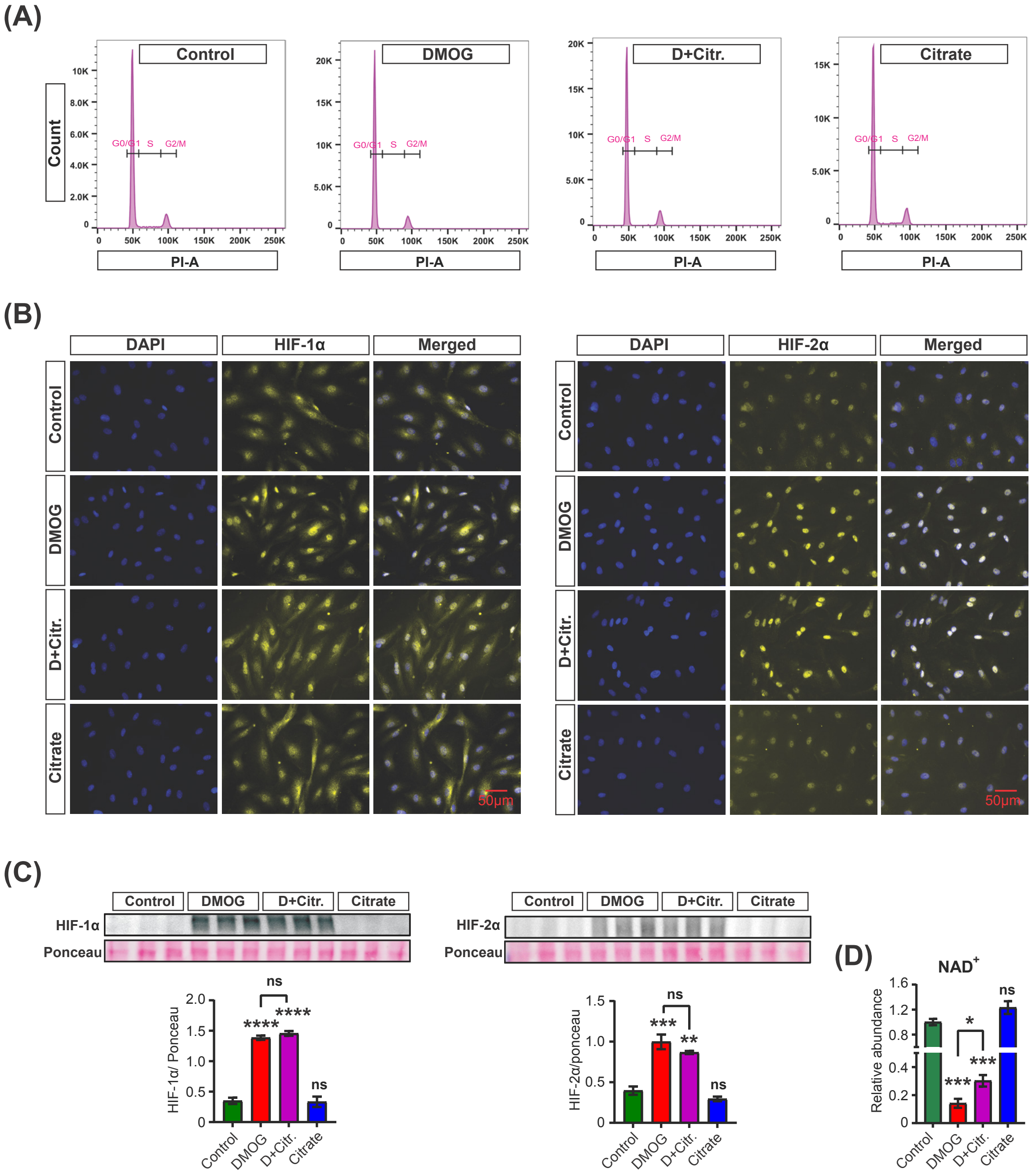
